# Design of two-stage multidrug chemotherapy schedules using replicator game dynamics

**DOI:** 10.1101/2024.07.16.603768

**Authors:** K. Stuckey, P.K. Newton

## Abstract

We use a replicator evolutionary game in conjunction with control theory to design a two-stage multidrug chemotherapy schedule where each stage has a specific design objective. In the first stage, we use optimal control theory that minimizes a cost function to design a *transfer orbit* which takes any initial tumor-cell frequency composition and steers it to a state-space region of three competing clonal subpopulations in which the three populations co-exist with a relatively equal abundance (high-entropy co-existence region). In the second stage, we use adaptive control with continuous monitoring of the subpopulation balance to design a *maintenance orbit* which keeps the subpopulations trapped in the favorable co-existence region to suppress the competitive release of a resistant cell population in order to avoid the onset of chemoresistance. Our controlled replicator dynamics model consists of a chemo-sensitive cell phenotype *S*, which is sensitive to both drugs, and two resistant cell phenotypes, *R*_1_ and *R*_2_, which are sensitive to drugs 1 and 2 respectively, but resistant to drug 2 and 1. The 3 × 3 payoff matrix used to define the fitness function associated with the interactions of the competing populations is a prisoner’s dilemma matrix which ensures that in the absence of chemotherapy, the *S* population (defectors) has higher fitness (reproductive prowess) than the two resistant cell populations, reflecting an inherent cost of resistance which our chemotherapy design methodology seeks to exploit. In our model, the two drugs *C*_1_ and *C*_2_ can act synergistically, additively, or antagonistically on the populations of cells as they compete and evolve under natural and artifical selection dynamics. Our model brings to light the inherent trade-offs between navigating to the maintenance orbit in minimal time vs. arriving there using the least total drug dose and also that the optimal balance of synergystic or antagonistic drug combinations depends the frequency balance of the populations of cells.

## I. INTRODUCTION

Multidrug chemotherapy dosing has been a mainstay in the treatment of cancers for over 50 years [1]. It has been widely appreciated that using toxic drug combinations that operate via orthogonal mechanisms on different clonal populations can be a powerful tool in both killing off different subpopulations of malignant cells as well as managing chemoresistance in a tumor [2]. As the drugs are administered and different subpopulations of cells are killed at different rates, the balance between different clonal tumor subpopulations changes, as does the subsequent underlying nature of their competition which is guided by frequency-dependent fitness principles [3]. Since different drug combinations apply different selection pressures on the subpopulations as they evolve, the frequency ratios of these subpopulations are continually changing. As they change, there is a need to manage the imbalance of the subpopulations in order to maintain intratumor cell competition so as to delay or avoid the evolution of chemoresistance which would ensue if one of the resistant phenotypes reached fixation [4]. An apt analogy is the recognized need to periodically rebalance financial portfolios as markets evolve, which, if neglected, would create large and risky imbalances among their components [5]. To avoid cancer recurrance due to chemotherapeutic resistance, the intelligent use of everchanging multidrug combinations is necessary, but in the absence of a quantitive multidrug dose-response model [6] in an evolving setting, this quickly develops into a combinatorial guessing game of high-stakes trial and error [7], particularly if many different drugs, designed to target different molecular mechanisms are involved [8]. Examples of efforts to address the combined challenge of controlling the evolutionary dynamics of the heterogeneous cell population constituting a tumor with combination therapies can be found in [9, 10].

As background to the methods developed in this paper, in [11] we laid out some general principles in guiding adaptive therapy that seeks to maintain a controllable stable tumor burden by capitalizing on the competitive interactions between drug-sensitive and drug-resistant subclones, which include: (i) identifying *frequencydependent cycles* of tumor evolution; (ii) realizing there are limitations to *evolutionary absorbing regions* reachable by the tumor; and (iii) leveraging the *rates of evolution* to determine optimal timing of drug sequencing. In addition, in [12] we highlighted the importance of leveraging both synergistic and antagonistic drug interactions, which has been developed and exploited more in the context of microbial populations to date [13, 14] than in tumor cell populations.

Here we develop a further important step within the framework of a three-component, two-drug replicator dynamics evolutionary game model, that of dividing the chemotherapy schedule into distinct stages with different design goals — here we focus on two-stage chemotherapy scheduling. The first, which we call the design of a *transfer orbit*, uses optimal control theory with a cost function that takes into account of the total chemodosage, total time, and final target in the three-component statespace. The goal of this first stage of chemo-scheduling is to take the initial three-component subpopulation profile and drive it to more favorable region of the state-space where the subpopulations compete on an equal footing (the high-entropy co-existance region). The second stage, in which we design a *maintenance orbit*, uses adaptive control theory to maintain a relatively equal balance of the three-subpopulations in order to supress the competitive release of a resistant population [15–17] while managing competition of the three cell phenotypes [18].

The idea of dividing a trajectory into components, each with its own design objective, is inspired by very succesful control theory strategies in the classical orbital and satellite dynamics literature, where the transfer orbit, called a *Hohmann transfer* [19–21], is commonly used to drive a satellite from one near-earth parking orbit to another. In our context, the transfer orbit plays the role of the Hohmann transfer, while the maintenance orbit (also called an evolutionary cycle [18]) plays the role of the parking orbit. Interestingly, in both the orbital mechanics context as well as the chemotherapy design context, the parking orbit must typically be maintained in a region where there is no asymptotically stable fixed point (for example, in a deep space region surrounding a libration point), so generally requires constant monitoring and adaptive controllers — reaching a target point and parking it there with no control input is generally not feasible. In the orbital mechanics context, the controllers are the rocket fuel propulsors which drive the rocket/satellite to a desired location, whereas in the context of tumor control, the controllers are the different chemo-dosing schedules [18] which we use to apply targeted selection pressure on the competing clonal populations.

While the advocacy of adaptive chemo-dosing schedules that effectively manage the intratumor cell competition (the maintenance orbit), instead of using open-loop control protocols (such as maximum tolerated dose (MTD) schedules) that can select for resistant populations, is not new [4], it is not yet widely accepted in the oncology community. This is partly due to the complexity of the required schedules, the difficulty in accurate real-time monitoring of subpopulation balances necessary to maintain closed-loop control protocols, and lack of clinical trial evidence on their efficacy. It is also hindered due to the prevailing dogma of using MTD to eliminate all cancer cells in order to cure the disease [22– By contrast, an adaptive maintenance schedule is designed primarily to manage the disease by keeping it in a favorable region of the state-space where the clonal subpopulations remain in balance. This approach has become more attractive with the increasing realization that, in many cases, MTD effectively selects for a resistant population [4, 25–28] by killing off the bulk of the sensitive cell population that had been supressing the growth of the resistant cells (known as competitive release). There is, however, a growing list of ongoing clinical trials that attempt to use adaptive scheduling concepts for different tumor types. What has not yet been addressed is the need for designing optimal chemoschedules that guide (transfer) the cell subpopulations from their initial state (determined via biopsy or high-resolution imaging) to the joining of a desirable maintenance orbit with quantifiable features, such as upper and lower bounds on eco-diversity balance and total chemo-dose per cycle, which, in principle, could be designed by the oncologist with the aid of a tailored interactive computational model [29, 30] along with multiregion biopsies, single-cell analysis, and liquid biopsies operating in parallel [31]. We consider the division of chemo-schedule design into different components, with different design objectives, joined together to create a computer-guided strategy, to be the next important step in understanding multidrug chemotherapy design and we flesh out the details in this paper in the context of a minimal three-component, two-drug controlled replicator dynamics evolutionary game [12, 18].

The quantitative metric we use to measure the ecological diversity [32] of the three malignant cell subpopulations (*S, R*_1_, *R*_2_), in the same spirit as used extensively in evolutionary ecology contexts [33], is the Shannon entropy (as opposed to the Simpson index [34]). We adopt it based on its simplicity in this context and interpretability in other problems where it has been applied [35]. With this measure, we view the state-space trajectories associated with our three-component evolutionary game as curves on an entropy surface in a trilinear state-space that are shaped and guided by our chemotherapy schedules. To maintain balanced competition among the subpopulations, we seek to maximize the entropy inside the triangular simplex. We do this while quantifying the important properties of appropriately designed maintenance orbits in high entropy regions of the triangular simplex. Since the maximum entropy point, (*S, R*_1_, *R*_2_) = (1*/*3, 1*/*3, 1*/*3) is not a fixed point of the dynamical system (i.e. an evolutionarily stable state (ESS)) we need to use active control theory to maintain it there or in nearby regions. For this we design (adaptive) chemotherapy schedules that trap the trajectory in a region of high-diversity where competition among the cell populations maintains a maximal balance (high entropy co-existence).

This defines our first task of identifying appropriate trapping zones between two constant (high) entropy contours on the entropy surface and to design chemotherapeutic schedules that maintain the trajectories in those zones perpetually on closed evolutionary cycles [18] — these are the *maintenance orbits*. To reach one of these orbits from any arbitrary initial state, we use optimal control theory (with a specified cost function) [36, 37] to design the optimal chemotherapy schedule that takes the trajectory from any specified initial state to the closed maintenance orbit inside the trapping zone — these are the *transfer orbits*. Once the two distinct legs of the full orbit are calculated and pieced together, the combination trajectory can then be obtained and its features analyzed. In particular, we focus on the inherent tradeoffs involved in arriving at the maintenance orbit in minimal time vs. arriving using minimal chemo-toxins. In addition, we highlight the importance of the interplay between the two-drug interactions (synergistic, additive, or antagonistic) and the initial subpopulation balance (i.e. the initial conditions of the governing dynamical system). We lay out this two-stage chemotherapy design process in the following sections and then discuss several further points regarding the model and possible extensions.

## II. MODEL

### A. Three-component two-drug interaction model

We use a three-component replicator dynamical system composed of three cell types, (*S, R*_1_, *R*_2_) ≡ (*x*_1_, *x*_2_, *x*_3_), which is controlled using two time-dependent chemotherapy functions, 0 ≤ *C*_1_(*t*) ≤ 1 and 0 ≤ *C*_2_(*t*) ≤ 1, with 0 ≤ *C*_1_(*t*) + *C*_2_(*t*) ≤ 1. The three component model includes one sensitive cell type, *S*, which is sensitive to both chemotherapy drugs and two resistant cell types, *R*_1_, which is sensitive to drug 1 but resistant to drug 2, and *R*_2_, which is sensitive to drug 2 but resistant to drug 1. In contrast to the single drug, single resistant cell type model that was explored in [18], the addition of a secondary chemotherapy drug and resistant cell type allows us to independently administer two drugs *C*_1_ and *C*_2_ and explore the important effects of drug interactions.

The dynamics of the system are governed by the replicator equations [38, 39]:

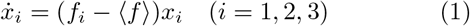

where *x*_*i*_ represents the frequency of cell population *i* = 1, 2, 3 and *x*_1_ + *x*_2_ + *x*_3_ = 1. The fitness function of each subpopulation, *f*_*i*_, is frequency dependent and given by:

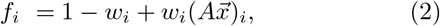

with the parameter 0 ≤ *w*_*i*_ ≤ 1 playing the role of a selection strength parameter. Note that when *w*_*i*_ = 0, *f*_*i*_ is constant, implying that selection plays no role in the resulting dynamics, whereas when *w*_*i*_ = 1, selection dominates the cell interactions. The interaction matrix *A* is the 3 × 3 payoff matrix that defines the game being played, and is given generically by

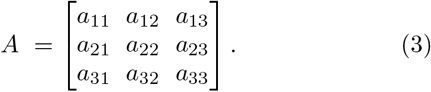

The average fitness of the tumor is given by ⟨*f*⟩ ≡ *f*_1_*x*_1_ + *f*_2_*x*_2_ + *f*_3_*x*_3_, thus the terms (*f*_*i*_−⟨*f*⟩) on the right of each equation, which is the deviation of each subpopulation fitness from the overall average fitness, represents the growth rate of that subpopulation. The equations constitute a cubic-nonlinear coupled system governing the three co-evolving subpopulations.

We embed the two time-dependent chemo-dosing functions (*C*_1_(*t*), *C*_2_(*t*)) into the parameter *w*_*i*_(*t*), 0 ≤ *w*_*i*_ ≤ 1, which determines the relative strength of selection as the subpopulations compete by setting:

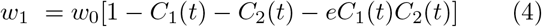

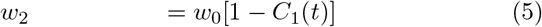

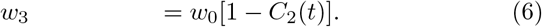

where *w*_0_ is a constant that scales time. If *w*_*i*_ ∼ 0, selection is weak and the evolutionary game being played does not have a large effect [40]. Conversely, if *w*_*i*_ ∼ 1, selection is strong in this case and the evolutionary game plays a larger role [41]. In our simulations, we take *w*_0_ = 0.1, which generally puts us in the weak selection regime usually considered appropriate for tumor cell competition. In the three cell, two drug model, eqn (4) gives the ability to create synergistic, additive, or antagonistic interactions by varying the parameter *e*. When *e* = 0 (or when either of the drugs is off), there is no cross-interaction acting on *x*_1_ (S), and the interaction is said to be additive — the combined effect of the two drugs is equivalent to the sum of the effects that the two drugs create independently. When *e >* 0, the selection parameter *w*_1_ is lowered more than it would otherwise be with both drugs on — this is called a synergistic interaction. When *e <* 0, the selection parameter *w*_1_ is lowered less than it would otherwise be with both drugs on — this is called an antagonistic interaction [12, 13].

The evolutionary game being played for our model is the prisoner’s dilemma (PD) game [39, 42]. For this, the entries of matrix *A* must satisfy the inequalities

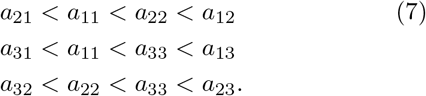

Under these conditions, we use the payoff matrix values

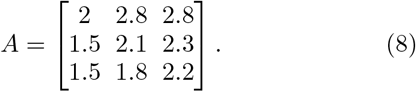

The PD game involves players that can choose between two actions: cooperation or defection. In the context of the PD game, the sensitive cells (*x*_1_) are the defectors and have a higher fitness level than the resistant cells types (*x*_2_ and *x*_3_) who are considered the cooperators. To see this most clearly, notice that the average of the entries in the top row of *A* is larger than that in the second row, which is larger than that in the third row. In classical (i.e. static) game theory, the players are thought to be making rational decisions with the goal of acquiring a higher payoff for themselves. In this case, the defection strategy for both players is a Nash equilibrium in which the average fitness is less than the cooperation strategy [43, 44]. The replicator dynamics model differs from classical game theory in the sense that the cells are not making strategic decisions but instead, their strategy is encoded in their reproductive prowess and strategies are heritable as a result of the replicator dynamics eqn (1). In this way, successful strategies (i.e. high reproductive prowess) are favored (via heritability) and emerge via natural selection dynamics [38, 45–4 The Nash equilibrium of the static game (defectors) is embedded into a nonlinear dynamical system with frequency dependent fitness, and the defector strategy emerges as an asymptotically stable state [48].

The PD game and replicator equations have been used when modeling cancer [50, 51] for the following reasons. First, they build in *survival of the fitter* by assigning higher frequency dependent fitness, and growth rates, to the sub-populations that are more fit than others (success breeds success). Second, they allow for the sensitive cells (with higher fitness due to the cost of resistance) to saturate the tumor in the absence of treatment (*C*_1_ = 0 and *C*_2_ = 0). Third, they ensure that this is a suboptimal state since in this state, the *S* cell population saturates the tumor and has a lower average fitness (⟨*f*⟩ = *a*_11_) than if one of the resistant cell types, *R*_1_ (⟨*f*⟩ = *a*_22_) or *R*_2_ (⟨*f*⟩ = *a*_33_), saturates the tumor. For added flexibibilty in our model, the two resistant populations, *R*_1_ and *R*_2_ have differing costs associated with two different resistance mechanisms, breaking the symmetry between the effects of *C*_1_ and *C*_2_ on these two subpopulations.

In order to understand key aspects of our model and our choice of parameters, we show trajectories of the equations in the (*S, R*_1_, *R*_2_) trilinear simplex in figure 1 for a range of chemotherapy parameter values. In the absence of chemotherapy (figure 1(a)), all trajectories converge to the *S* corner which is an evolutionary stable state (ESS), slowing down as the trajectory approaches the corner. The background color field indicates the instantaneous velocity to the *S* corner. Figures 1(b),(c),(d) lay out what happens as the *C*_2_ parameter is increased. Below threshold (figure 1(b)), the *S* corner remains the ESS. Above threshold (figure 1(c)), all trajectories converge to the *R*_1_ corner which becomes the new ESS. This is due to the fact that *C*_2_ acts on both the *S* and the *R*_2_ populations, releasing *R*_1_ from the competition (i.e. competitive release [15, 52, 53]). Figure (figure 1(d)) confirms that for even higher values of *C*_2_, the *R*_1_ corner is the ESS. Likewise, figures 1(e),(f),(g) show the competitive release of the *R*_2_ population as the *C*_1_ parameter is increased above a threshold value. In figure 1(h) we show one mixed case with parameter values *C*_1_ = 0.3, *C*_2_ = 0.3 (additive drug interaction parameter *e* = 0). For this, there is a separatrix curve which separates the basin boundary for convergence to the *S* corner vs. the *R*_1_ corner.

**FIG. 1.**
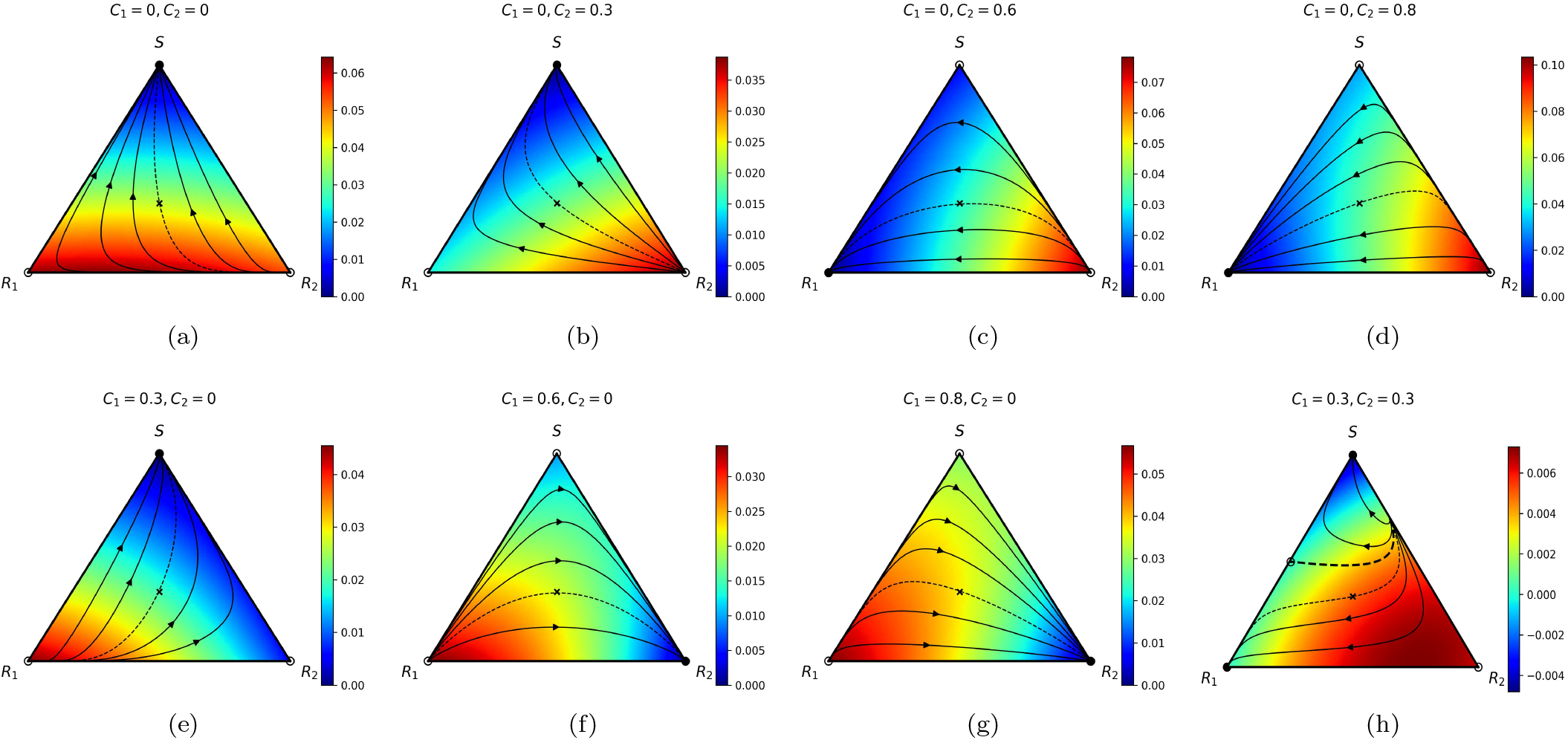
Trajectories and velocity heat maps to *S* corner for different (*C*_1_, *C*_2_) parameter values. (a)(*C*_1_, *C*_2_) = (0, 0). With no chemotherapy, all trajectories converge to *S* which is an ESS; (b)(*C*_1_, *C*_2_) = (0, 0.3). All trajectories converge to *S*, which is an ESS; (c) (*C*_1_, *C*_2_) = (0, 0.6). All trajectories converge to *R*_1_ which is an ESS; (d) (*C*_1_, *C*_2_) = (0, 0.8). All trajectories converge to *R*_1_ which is an ESS;(e)(*C*_1_, *C*_2_) = (0.3, 0). All trajectories converge to *S* which is an ESS; (f)(*C*_1_, *C*_2_) = (0.6, 0).All trajectories converge to *R*_2_ which is an ESS; (g)(*C*_1_, *C*_2_) = (0.8, 0). All trajectories converge to *R*_2_ which is an ESS; (h) (*C*_1_, *C*_2_) = (0.3, 0.3). Mixed state, with trajectories starting above the separatrix (dashed curve) converging to *S*, while trajectories starting below the dashed curve converging to *R*_1_.

### B. Measuring population diversity

As a measure of diversity of the three populations, we adopt the Shannon entropy criterion [34, 35], given by the formula:

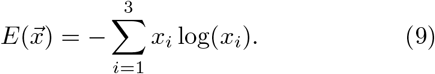

with *x*_1_ +*x*_2_ +*x*_3_ = 1, we have upper and lower bounds:

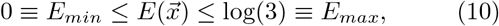

where we refer to *E*_*max*_ has the maximal co-existence point ((*x*_1_, *x*_2_, *x*_3_) = (1*/*3, 1*/*3, 1*/*3)). Note that this (central) point is not a fixed point of the dynamical system (1). We scale *E* by *E*_*max*_ and define a unit normalized Shannon entropy:

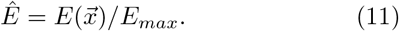

In figure 2 we show contour values of the entropy surface with maximium point located at 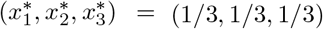. In this figure we also show nine representitive initial conditions, and paths of steepest ascent to *E*_*max*_ (gradient paths), perpendicular to constant entropy contours. These are the most direct paths from their initial points to the entropy peak (although not geodesic curves on the surface [54]). Which of these paths can be created using chemotherapy scheduling with functions (*C*_1_(*t*), *C*_2_(*t*)) will be described later.

**FIG. 2.**
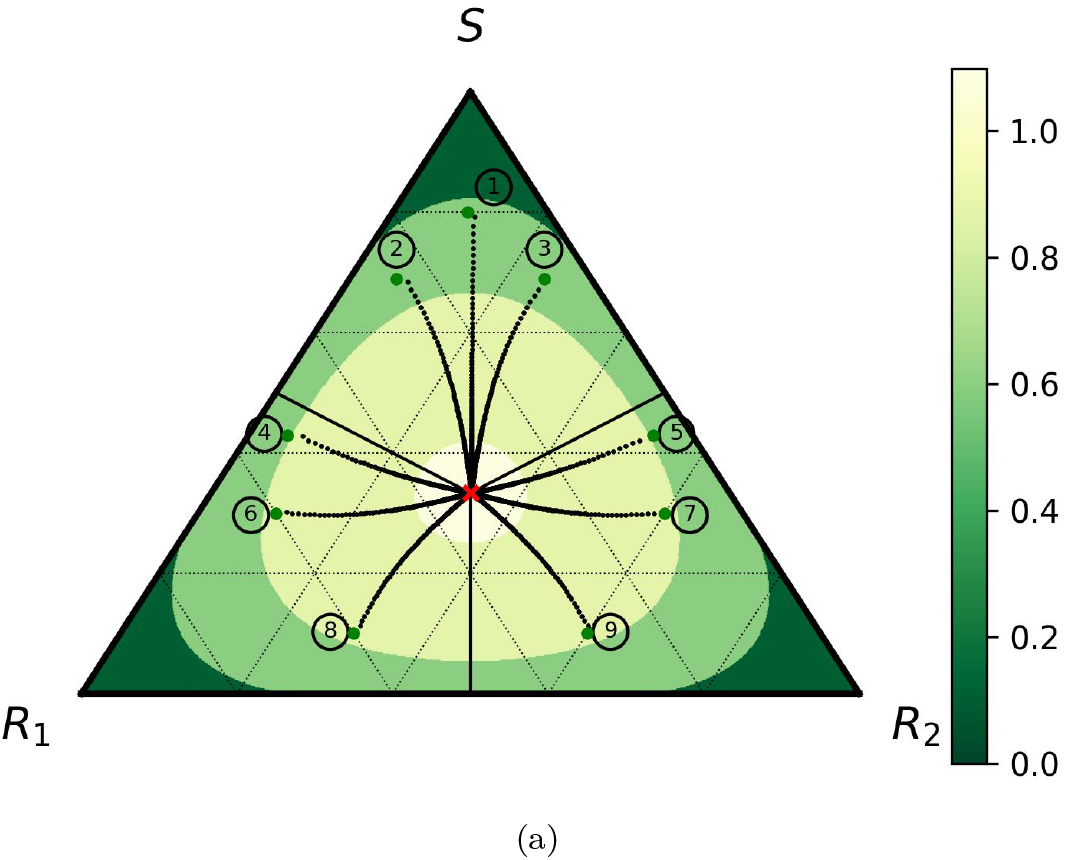
Steepest ascent paths on the entropy surface. Shown in color are the entropy levels 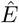 with color bar. Also shown are nine representative steepest ascent paths to *E*_*max*_. These paths represent the most direct path on the surface to *E*_*max*_ located at (*x*_1_, *x*_2_, *x*_3_) = (1*/*3, 1*/*3, 1*/*3).

## III. RESULTS

Next, we use the time-dependent controllers (*C*_1_(*t*), *C*_2_(*t*)) to design paths on the entropy surface.

### A. Constructing maintenance orbits via adaptive control

In figure 3 we lay out the procedure of constructing maintenance orbits. Figure 3(a) shows a continuous family of closed orbits of the dynamical system surrounding *E*_*max*_ at the center of the simplex. As noted in the caption, these curves are obtained by overlaying families of constant chemotherapy parameter curves (such as those shown in figure 1) ontop of each other, for different *C*_1_ and *C*_2_ values. We have included dashed lines of the same coloring scheme which show trajectories that intersect *E*_*max*_, i.e the point where the Shannon entropy function has a global maximum. The lowest entropy states are the corner points of the triangle given by 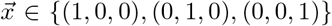. At these points, two of the subpopulations have gone extinct and the population is made entirely of one type of cell. Note that the corner points are fixed points of (1), but *E*_*max*_ is not.

**FIG. 3.**
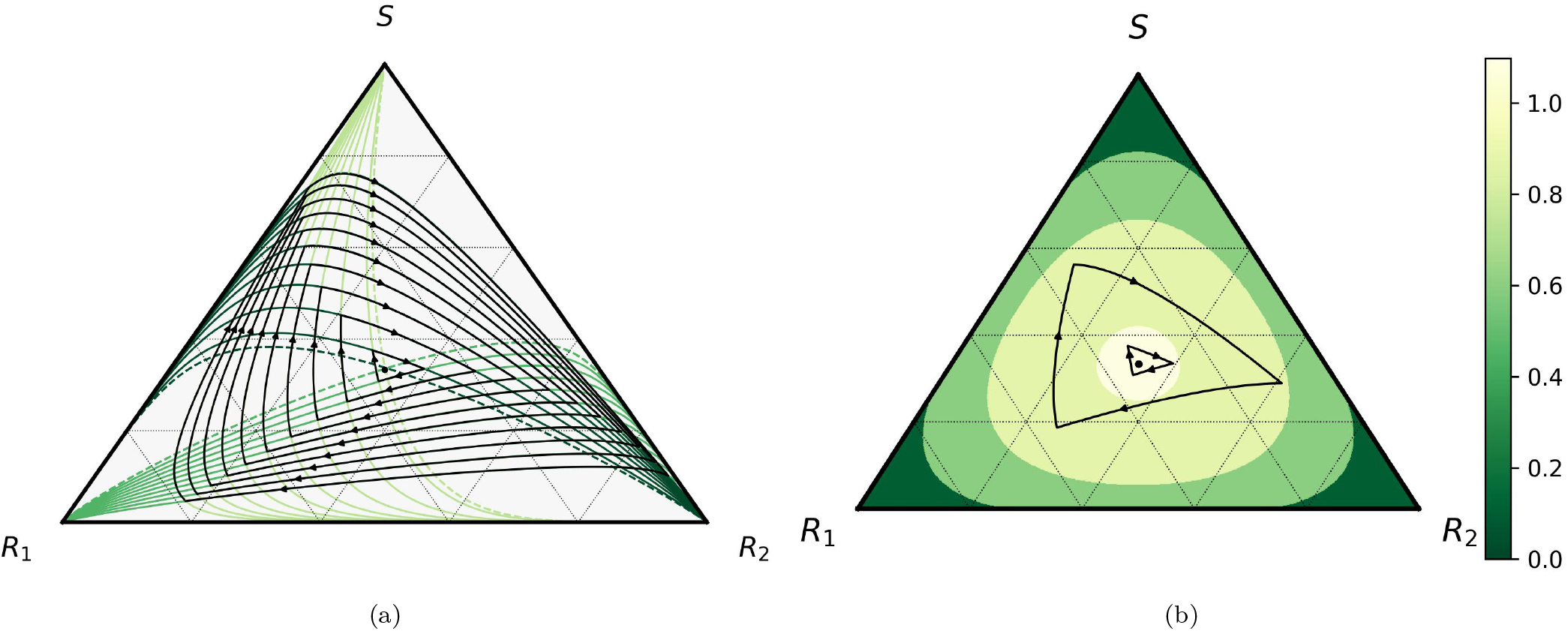
Family of closed evolutionary cycles in the *S, R*_1_, *R*_2_ triangular plane. (a) With no chemotherapy, represented by trajectories in light green, the tumor saturates to the *S* corner. When *C*_1_ = 0 and *C*_2_ = 0.8, represented by trajectories in green, competitive release of the resistant population *R*_1_ drives all trajectories to the *R*_1_ corner. When *C*_1_ = 0.8 and *C*_2_ = 0, represented by trajectories in dark green, competitive release of the resistant population *R*_2_ drives all trajectories to the *R*_2_ corner. The dashed lines represent trajectories that intersect the maximum entropy point where all three subpopulations are equal, 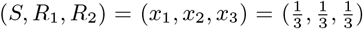. Overlaying the solution trajectories for the three constant chemotherapy profiles creates closed periodic orbits with a variety of enclosed areas. (b) Contours of constant entropy are shown with the lowest entropy contour shown in dark-green and highest entropy contour shown in white. We can design the adaptive cycle to be restricted between any two contours of constant entropy which traps the closed maintenance cycle between upper and lower entropy bounds.

To dynamically construct one of the closed maintenance orbits in the family, it is necessary to compute the times where two of the trajectories cross (i.e. the three corners of the closed loop), say *t* = *T*_1_ *< T*_2_ *< T*_3_ at respective corners. This defines a piecewise constant maintenance chemotherapy schedule for that closed loop:

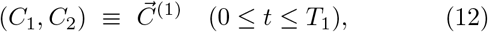

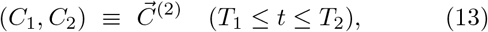

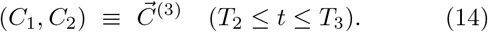

Each of the closed loops shown in figure 3(a) lies on the entropy surface shown in figure 3(b). Note, however, that the a closed loop is not a constant entropy curve. The dark green represents a contour where the entropy is close to zero. As we move towards the center of the triangle, we have a closed contour that contains the maximum entropy point, labeled in white. In this figure, we see that we can design a maintenance orbit that is trapped between contours of constant entropy. Because of this, we can construct maintenance orbits that trap the entropy level below and above entropy values of our choosing.

Figures 4 and 5 show the properties of the full family of maintenance orbits as a function of the area they enclose (figure 4), and the time to complete one cycle (figure 5). Figure 4(a) shows that the total chemo-dosages used increase as a function of the area enclosed, while figure 4(b) shows the perimeter of the closed loop as a function of area, which is close to that of an equiliateral triangle. In figures 5(a),(b),(c) we show properties of the maintenance cycles as a function of time to complete the cycle. The area enclosed by a cycle increases (roughly quadratically) with the time to complete the cycle (figure 5(a)); the total chemo-dose used increases linearly with time (figure 5(b)); and the perimeter of the cycle increases (roughly linearly) with time to complete the cycle (figure 5(c)).

**FIG. 4.**
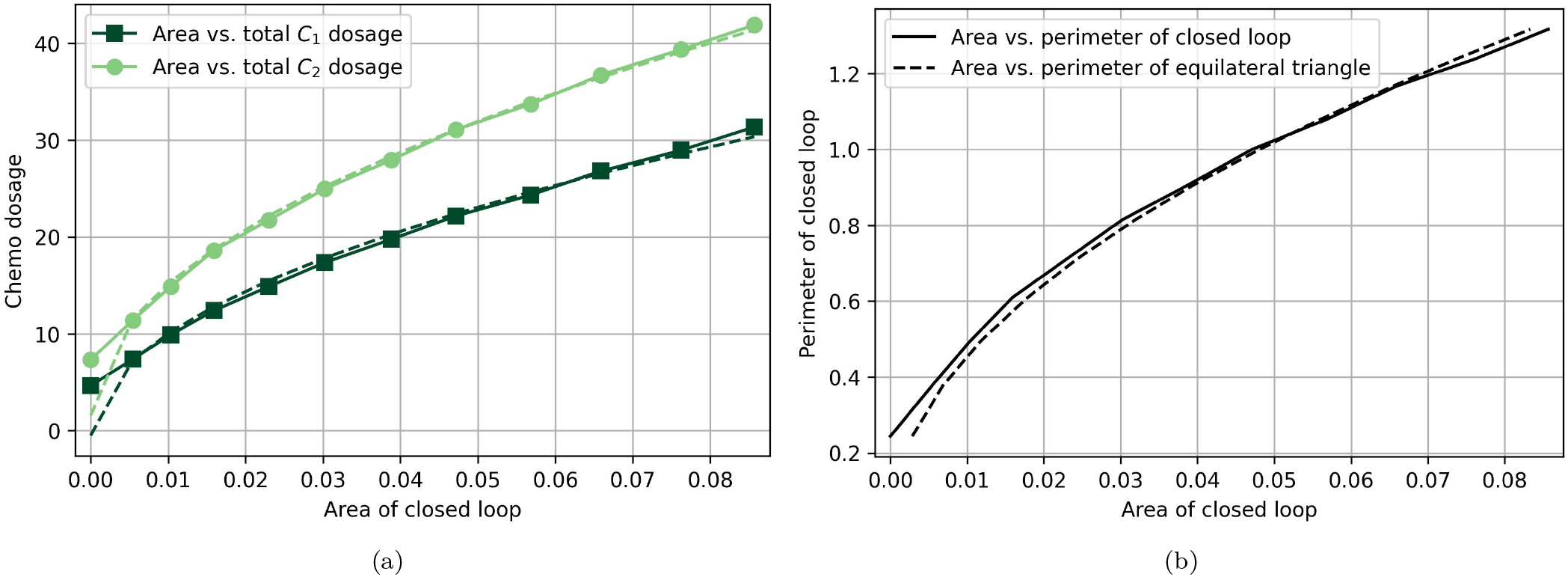
Properties of evolutionary cycles as a function of area. (a) Total chemotherapy dosage administered versus the enclosed areas of the maintenance cycles. (b) Perimeter versus the enclosed areas of the maintenance cycles.

**FIG. 5.**
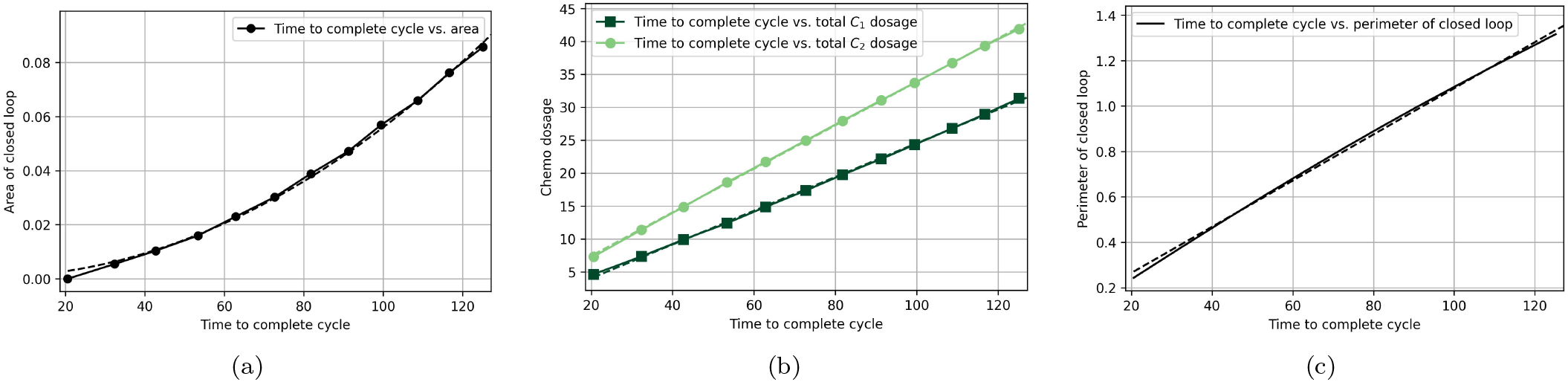
Properties of the maintenance cycles as a function of time. (a) Enclosed areas of the maintenace cycles versus the time to complete the cycles. (b) Total chemotherapy dosage administered versus the time to complete the cycles. (c) Perimeter of the maintenance cycles versus the time to complete the cycles.

### B. Constructing transfer orbits via optimal control

We turn next to the construction of the transfer orbits using optimal control [37, 55–57]. We briefly introduce an optimal control framework with the goal of minimizing a chosen cost function. The controller in this model is a vector of the two administered chemotherapy drugs, 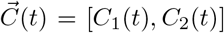 which are embedded in the fitness parameters *w*_*i*_ via eqns (4)-(6). We would like to find an admissible controller which causes the system to follow a solution trajectory that minimizes a desired performance measure, or cost function:

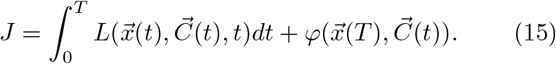

In particular, when we say that a controller is optimal, it indicates that

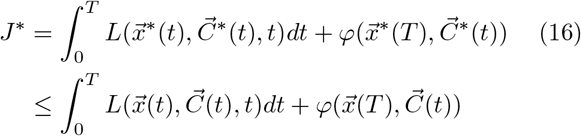

for all 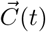 and corresponding solution trajectory 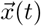. Here, we denote the optimal control schedule as 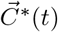 and the optimal trajectory as 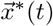. This inequality ensures that the optimal control and trajectory give us a global minimum of the performance measure *J*. We define the total dose delivered in time *t*, 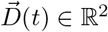, as

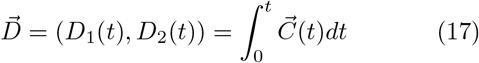

and thus,

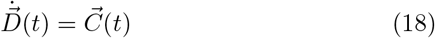

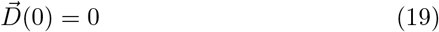

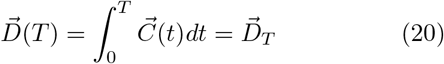

where *T* denotes the final time in which we implement control. We enforce constraints on the controllers such that

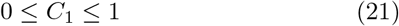

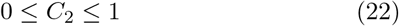

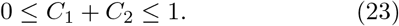

Using the above formulation, we compute trajectories from any initial state to a chosen maintenance orbit that closely surrounds the maximum entropy point by minimizing the cost function (15). The first term in (15) is called the running cost, while the second is the terminal cost. We identified a contour of constant entropy that intersects the chosen maintenance orbit at six intersection points and from eight initial conditions, labeled 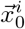, we optimally drive the trajectory to the closest intersection point, labeled 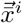, at some (undetermined) final time *T*. This is shown in figure 6. The cost function is made up of our chemotherapy dosing functions, target points on the maintenance orbit, and end time *T* :

**FIG. 6.**
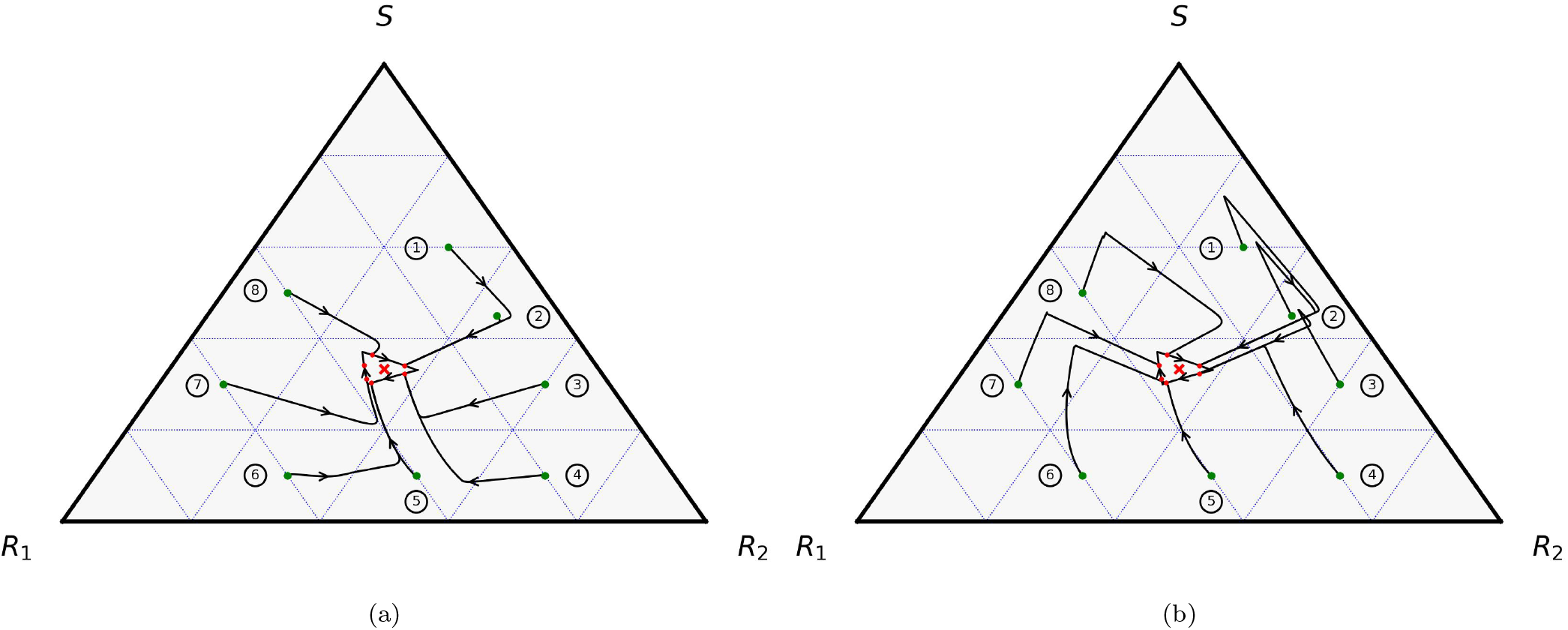
Transfer orbit design using optimal control. The optimal solution trajectory found for eight initial conditions when (a) *α* = 1, *β* = 0, *γ* = 10^−5^ and (b) *α* = 1, *β* = 0.05, *γ* = 10^−5^ when minimizing the distance from the solution trajectory to the chosen parking orbit. When the trajectory intersects the closed maintenace orbit, the controllers switch to a piecewise constant chemotherapy dosage schedule.

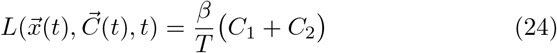

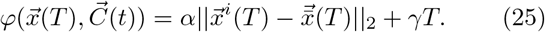

The running cost measures the total dosage through time *T*. The terminal cost is made up of two parts- the first term measures the final distance to the maintenance orbit, while the second term measures the total time to traverse from the initial point to the maintenance orbit. We have introduced three parameters into our optimal control model, *α, β* and *γ*, that can be altered to explore the effects that different terms have on the transfer orbits.

All solutions to this problem were computed using GEKKO, which is a Python package for machine learning and optimization of mixed-integer and differential algebraic equations. Documentation for this package can be found at https://gekko.readthedocs.io/en/latest/.

### C. Target hitting transfer trajectories

The first sub-problem we address is a Mayer problem (target only) in which we minimize the final distance between the solution trajectory and the chosen maintenance orbit at time *T*. In this case, we set the parameter *β* to zero and thus, the cost function to be minimized is given as

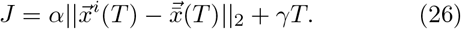

Note that, when the parameters are chosen this way, we are imposing a terminal criterion only, with no running cost. In Figure 6(a), we compute the optimal solution trajectory from eight initial conditions with *α* = 1 and *γ* = 1*e* − 5. Combining optimal control framework with the maintenance orbits found in the previous section, we find the optimal chemotherapy dosage schedule with corresponding trajectories that take these initial conditions towards the maintenace orbit region. These are shown in figure 7. In figure 6(a), we have highlighted a chosen closed maintenace orbit that tightly encloses the maximum entropy point (i.e. the upper and lower bounds on the entropy values along this orbit do not vary widely). When the transfer trajectory intersects this maintenance orbit, we switch to a piecewise constant chemotherapy schedule. This keeps the solution trajectory orbiting around the maximum entropy point indefinitely. In Figure 7, we show the chemotherapy dosage schedules which correspond to the same initial conditions seen in Figure 6(a). Here, the chemotherapy schedule before the thick black vertical line indicates the optimal schedule until it reaches the closed maintenance orbit. After the thick black line, we use the piecewise constant schedule used to maintain the orbit around the closed loop. In Table I we show the time (*T*) to traverse the transfer orbit to the maintenance orbit for each of the eight initial conditions shown. In general, the three inital conditions in the upper region of the triangle (marked as 1, 2, 8) arrive at the maintenace orbit points more quickly than those in the lower part of the triangle.

**FIG. 7.**
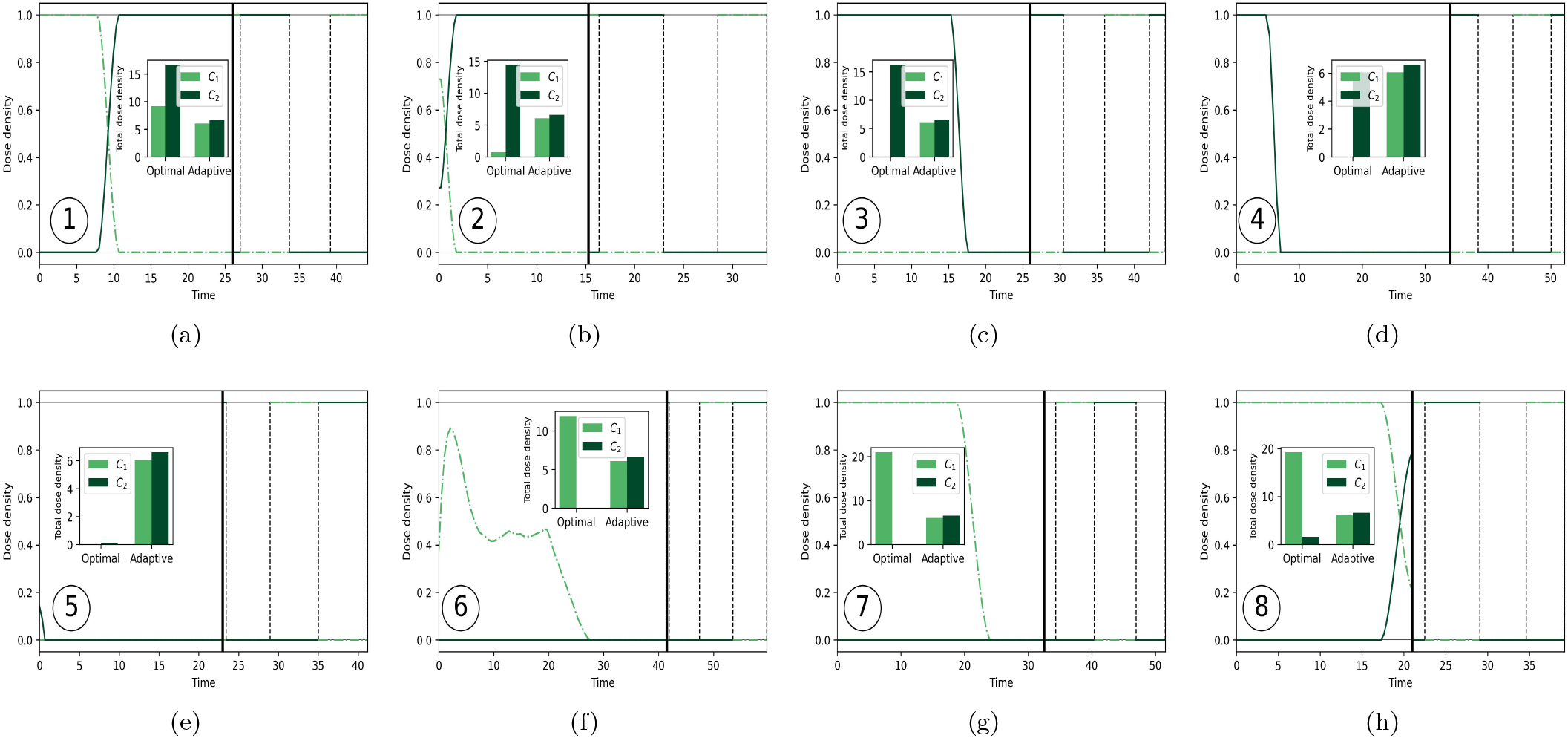
Chemotherapy dosage schedules for corresponding eight initial conditions in Figure 6(a). These plots correspond to the eight initial conditions used in Figure 6(a). The dark green dashed dotted lines indicate the dose density of the chemotherapy drug *C*_1_ while the light green lines indicate the dose density of the chemotherapy drug *C*_2_. The thick black line shows the time in which we switch from the optimal chemotherapy schedule to the closed periodic orbit. Here, we show the maintenance schedule for one cycle. Also included in these plots is the total dose density of each chemotherapy drug where we compare the amount of chemotherapy drugs used in the optimal (transfer) versus maintenance schedules. The initial conditions used are (a) *x*_0_ = [0.6, 0.1, 0.3], (b) *x*_0_ = [0.45, 0.1, 0.45], (c) *x*_0_ = [0.3, 0.1, 0.6], (d) *x*_0_ = [0.1, 0.2, 0.7], (e) *x*_0_ = [0.1, 0.4, 0.5], (f) *x*_0_ = [0.1, 0.6, 0.3], (g) *x*_0_ = [0.3, 0.6, 0.1] and (h) *x*_0_ = [0.5, 0.4, 0.1].

**TABLE I.**
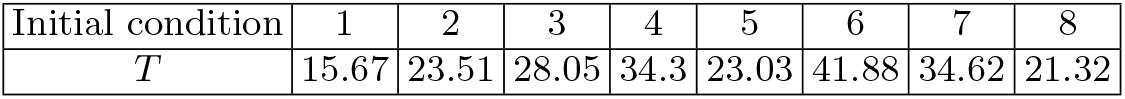
Transfer orbit time with *α* = 1, *β* = 0, *γ* = 10^−5^.

| Initial condition | 1 | 2 | 3 | 4 | 5 | 6 | 7 | 8 |
| --- | --- | --- | --- | --- | --- | --- | --- | --- |
| $T$ | 15.67 | 23.51 | 28.05 | 34.3 | 23.03 | 41.88 | 34.62 | 21.32 |

### D. Minimum chemo-dose transfer trajectories

In the secondary subproblem, in addition to minimizing the distance between the solution trajectory and the chosen parking orbit at time *T*, we consider the total administered chemotherapy dosage. In this case, we set the parameters *α* = 1, *β* = 0.05 and *γ* = 1*e* − 5 and thus, the cost function to be minimized is given as

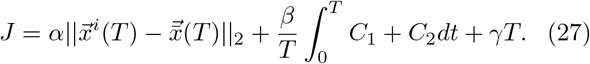

In Figure 6(b), we again use the same eight initial conditions that were used in the previous subproblem and find the optimal chemotherapy dosage schedule that minimizes our chosen cost function. Figure 8 shows the chemotherapy dosage schedules for these eight initial conditions.

**FIG. 8.**
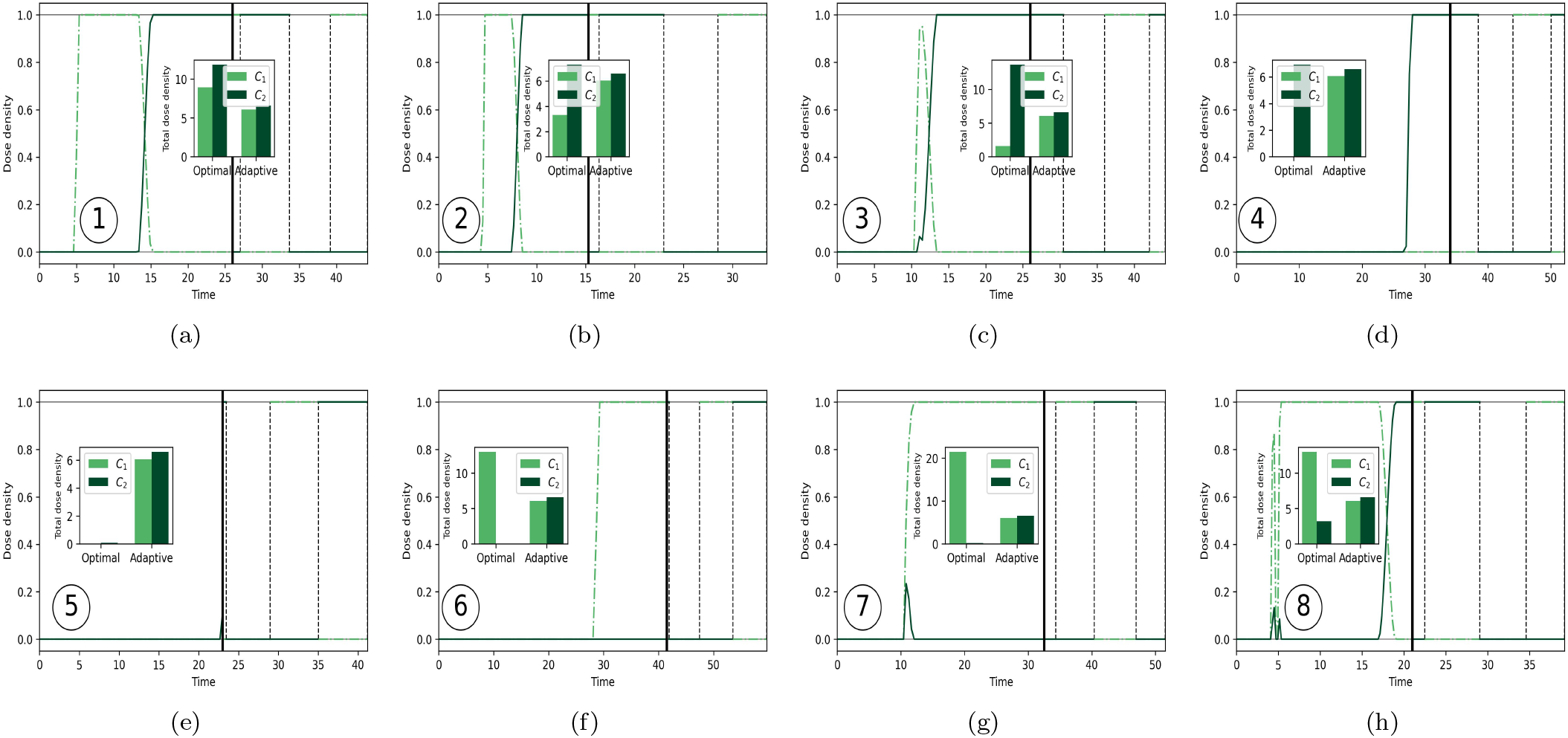
Chemotherapy dosage schedules for corresponding eight initial conditions in Figure 6(b). The dark green dashed dotted lines indicate the dose density of the chemotherapy drug *C*_1_ while the light green lines indicate the dose density of the chemotherapy drug *C*_2_. The thick black line shows the time in which we switch from the optimal (transfer) chemotherapy schedule to the closed periodic maintenance orbit. Here, we show the maintenance schedule for one cycle. Also included in these plots is the total dose density of each chemotherapy drug where we compare the amount of chemotherapy drugs used in the optimal (transfer) versus maintenance schedules. The initial conditions used were (a) *x*_0_ = [0.6, 0.1, 0.3], (b) *x*_0_ = [0.45, 0.1, 0.45], (c) *x*_0_ = [0.3, 0.1, 0.6], (d) *x*_0_ = [0.1, 0.2, 0.7], (e) *x*_0_ = [0.1, 0.4, 0.5], (f) *x*_0_ = [0.1, 0.6, 0.3], (g) *x*_0_ = [0.3, 0.6, 0.1] and (h) *x*_0_ = [0.5, 0.4, 0.1].

In figure 9 we show the average dose density used to obtain the trajectories in figures 6(a),(b). Tables I and II show the time needed to traverse the transfer orbit for each of the two cases. Table II shows the time (*T*) to the maintenace orbit, which generally are much longer than those shown in Table I.

**FIG. 9.**
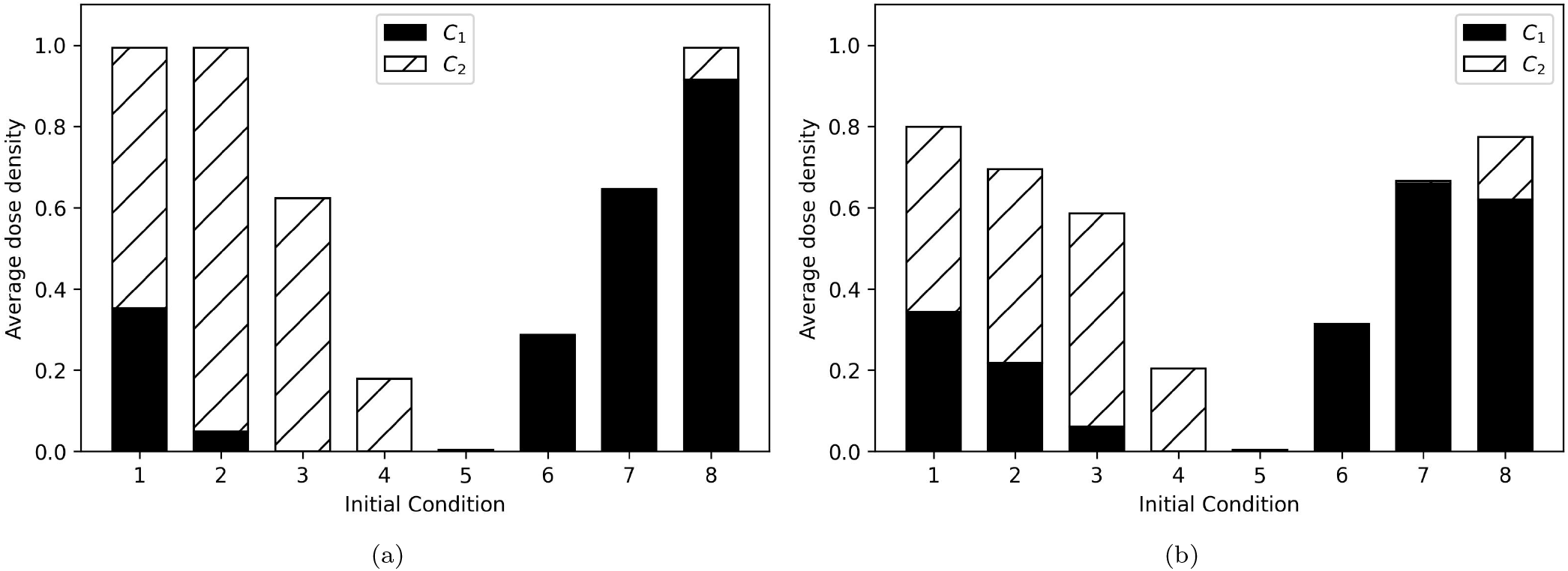
Average dose density for trajectories in Figure 6(a) and Figure 6(b). Average dose density when (a) *α* = 1, *β* = 0, *γ* = 10^−5^ and (b) *α* = 1, *β* = 0.05, *γ* = 10^−5^.

**TABLE II.** Transfer orbit time with *α* = 1, *β* = 0.05, *γ* = 10^−5^.

| Initial condition | 1 | 2 | 3 | 4 | 5 | 6 | 7 | 8 |
| --- | --- | --- | --- | --- | --- | --- | --- | --- |
| $T$ | 42.96 | 33 | 50.62 | 38.61 | 23.01 | 42.15 | 36.53 | 47.13 |

The trade-off between minimizing the time to the maintenace orbit vs. minimizing the total chemo-dosage (dose density) is shown clearly in figure 9. Minimizing the time to the maintenace orbit (*β* = 0) comes at the cost, shown in figure 9(a) of having to use a higher dose density. On the other hand, when we include the running cost into the cost function (*β* ≠ 0), as shown in figure 9(b), we reduce the total chemo-dose delivered, but from Table II, it takes more time to arrive at the target. This makes sense in light of the analogy between this problem and the Hohmann transfer and parking orbits from satellite/orbital mechanics: it generally takes more *fuel* (chemo-toxins) to arrive at a destination more quickly — there is a trade-off between fuel consumption and tranfer time.

### E. Multi-drug synergy and antagonism

In figure 10 we explore the effects of drug synergy (*e >* 0), and antagonism (*e <* 0), again for nine distributed initial conditions. From the upper part of the triangular simplex shown in 10(a) (initial conditions 1, 2, 3), there are clear differences in the optimal trajectories heading to the maximum entropy point depending on whether the drug interactions are synergystic, additive, or antagonistic. Generally speaking, the synergistic trajectories (*e* = 0.5) take a more direct path down to the center point than the additive or antagonistic paths. They also use more chemotherapy, as shown in 10(b),(c),(g). Initial conditions 4 − 9 in the lower part of the triangular simplex (10(a)) follow the same trajectories regardless of the value of the parameter *e*. One can understand why by looking at the total dose density histograms in figure 10(d),(e),(f),(h),(i),(j) for these initial conditions — in each case, either *C*_1_ or *C*_2_ is turned off and the optimal trajectory is guided by a single drug. In those cases, from eqn (4) one can see that there is no cross-term in the selection pressure parameter, so there is no opportunity for synergy or antagonism. It is interesting and noteworthy that the effect of synergy vs. antagonism depends very much on the initial location point in the triangular simplex, i.e. the subpopulation balance is part of the determination of whether or not the drugs interact synergistically or antagonistically in an co-evolving setting.

**FIG. 10.**
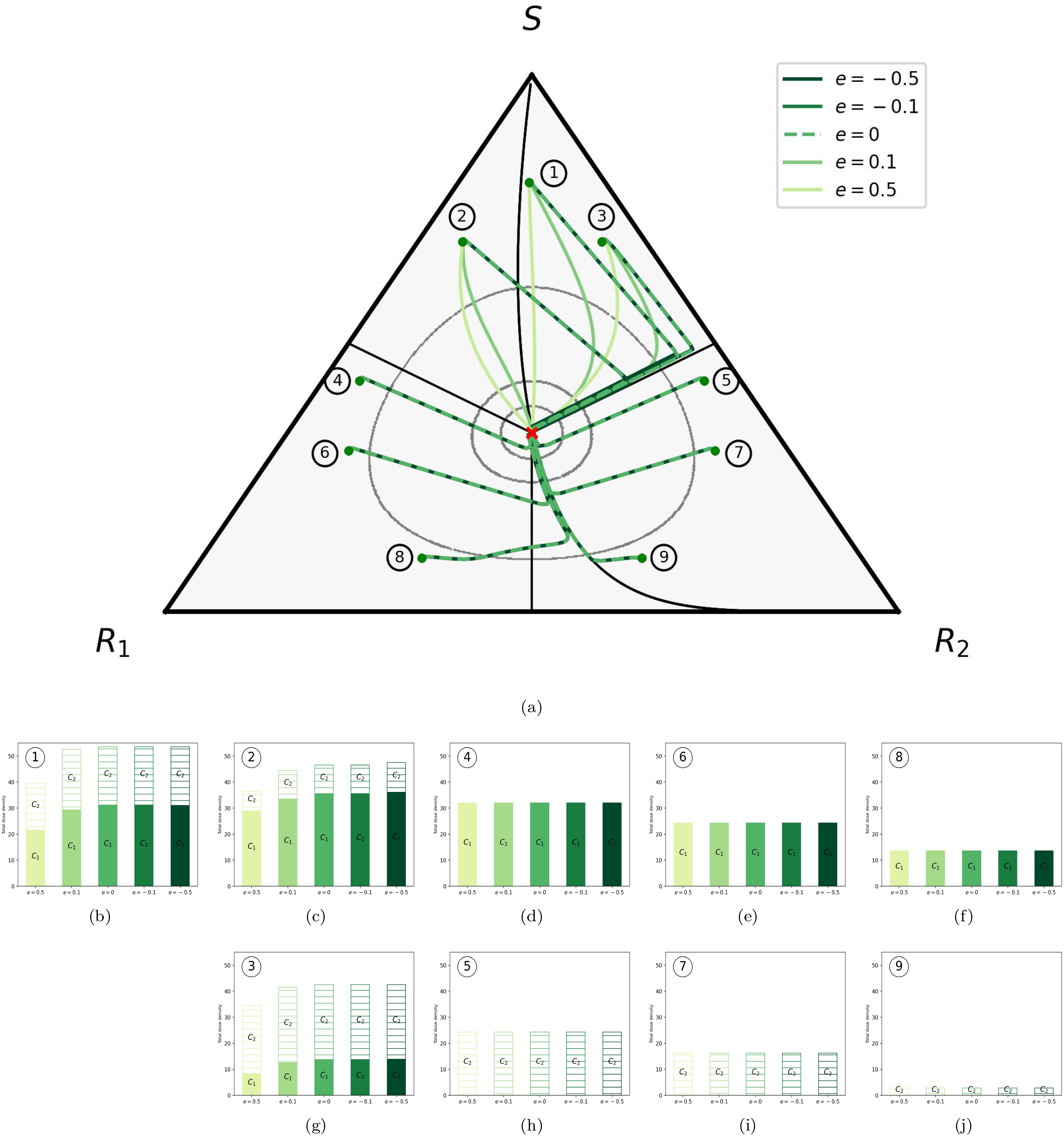
The role of synergy (*e >* 0) and antagonism (*e <* 0) within eight regions of the triangular phase plane. (a) The regions are divided by the symmetry lines of the triangle and the three trajectories that intersect the maximum entropy point. The optimal trajectory was found for an initial condition within each of these regions. The synergistic (solid, lighter green), additive (dashed green) and antagonistic effects (solid, darker green) were compared. (b) Panels of the total dose density for the nine initial conditions. Here, the darker color indicates the total dose density of chemotherapy drug *C*_1_ and the lighter color indicates the total dose density of the chemotherapy drug *C*_2_.

As a final point, we focus on the initial condition 1 in figure 10(a) and compare the resulting path to that of the corresponding gradient ascent path 1 from figure 2, noting they are quite similar. We ask, is the optimal dose density that created this path, shown in figure 11(a), similar to a dose density that would create the gradient ascent path? The two dose densities are compared in figure 11(b) with synergystic drug interactions (*e* = 0.5) — the two are very similar. Our conclusion is that the optimal path from initial condition 1, with synergistic drug interactions, as very nearly the gradient ascent path to the center point. The two paths are compared in 11(c). For all other initial conditions, the optimal paths, and the gradient ascent paths are quite different [54] — it is generally not possible to follow gradient ascent paths using chemo-dose schedules of the kind developed in this manuscript.

**FIG. 11.**
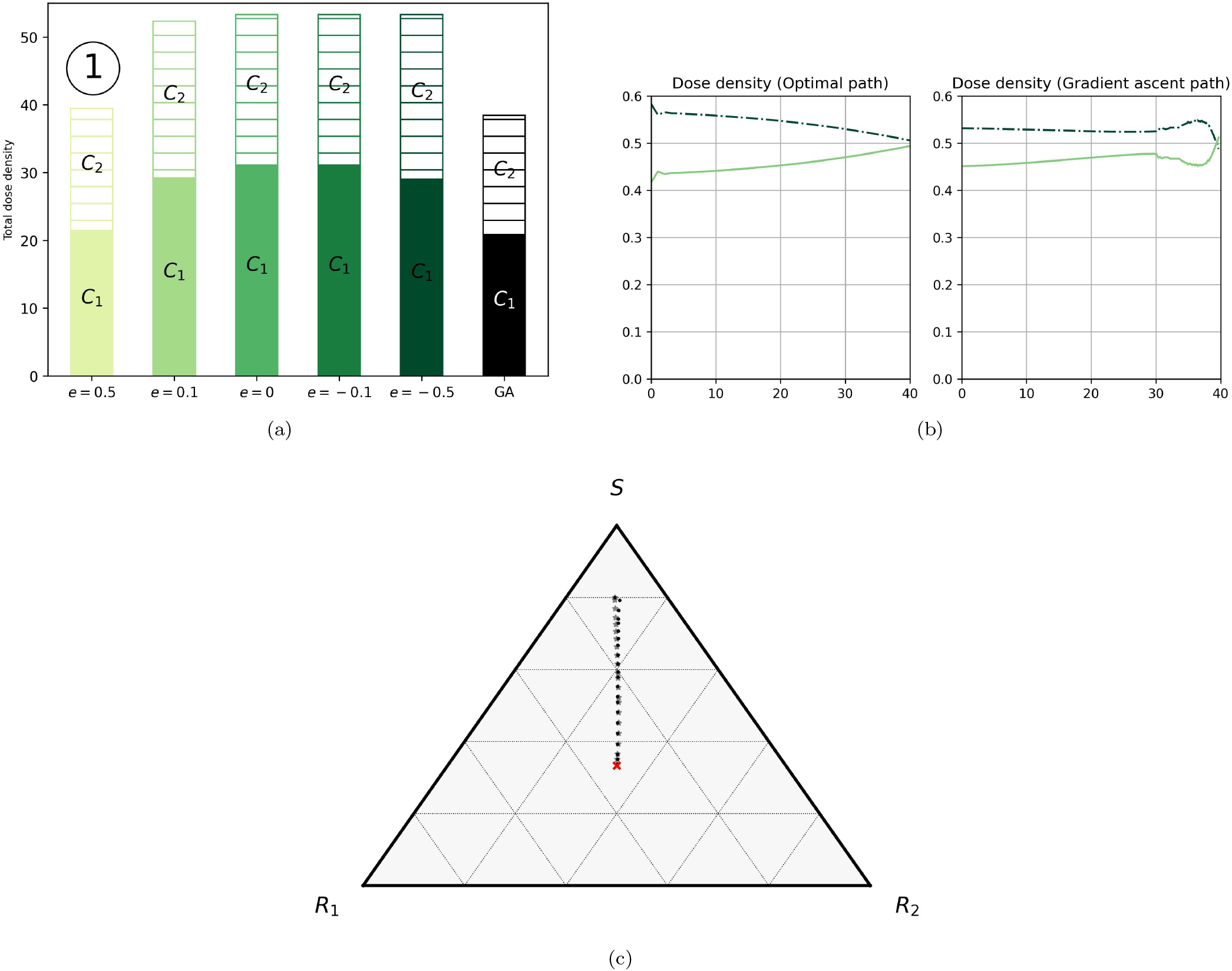
Comparison between optimal (transfer) solutions and gradient ascent paths. (a) Total chemotherapy dosage used for initial condition 1 to reach the maximum entropy point for the Mayer problem in 10(a). (b) Chemotherapy schedule corresponding to optimal trajectory for initial condition 1 in 10(a) (*e* = 0.5) and the near gradient ascent path. (c) Comparison of optimal trajectory for initial condition 1 (*e* = 0.5) with near gradient ascent path produced by a chemo-schedule in (b).

## IV. DISCUSSION

The concept of dividing the chemotherapy scheduling into two distict stages appears to be a novel concept, but is natural given the realization that each of the two stages has a distinct objective. In the transfer stage, the goal is to drive the subpopulation balance to a high-entropy region of the trilinear simplex where the subpopulations compete on equal footing with each other without allowing any to reach fixation. This ensures that neither of the resistant populations will saturate the tumor. For this, optimal control theory is an ideal design tool that clearly highlights the trade-offs between arriving quickly vs. arriving without using high levels of chemo-toxins. Once in the desirable region of the state-space, switching to a maintenance orbit using adaptive control with monitoring and steering is more natural in a setting in which the subpopulations are evolving via natural selection dynamics — optimizing in these settings is generally not productive.

The role of synergy vs. antagonism for drug interactions has been studied extensively in static settings [58, 59], but much less in settings in which the populations are co-evolving [12, 13, 60]. Drug interactions, in general, are much better understood in evolving microbial populations than in tumor cell populations. What had not been emphasized or perhaps fully appreciated previously in these works is the important role that the baseline population balance plays in the effects of drug interactions, and this could and should be further explored.

For longer term maintenance of the subpopulation balance, particularly in a stochastic environment [61], we expect that a third stage of the chemotherapy schedule would be needed to counteract that stochastic drift pushing the maintenance orbit off its closed loop [53, 62]. This third stage could be called the *stochastic adjustment* stage and has only been explored briefly in [62]. Crucial in the accurate calculation of these multi-stage chemoschedules is the importance of accurate monitoring of the population balances in these contexts [63]. While this is currently a clinical challenge, our view is that methods for accurately monitering of cellular subpopulations in an evolving tumor [31] will continue to improve over time making state-space optimal and adaptive control theory ideas increasingly feasible.

## ACKNOWLEDGMENTS

We gratefully acknowledge support from the Army Research Office MURI Award #W911NF1910269 (2019-2024).

